# Metabolic signatures of ferritin and TDP-43 co-pathology provide a mechanistic basis for stratified therapeutic approaches in ALS

**DOI:** 10.64898/2026.03.13.711539

**Authors:** Holly Spence, Fiona L. Read, Fergal M. Waldron, Jenna M. Gregory

## Abstract

**Background:** ALS is increasingly recognized as a biologically heterogeneous disease in which several molecular and pathological mechanisms converge on a similar clinical phenotype. One of these molecular markers is ferritin accumulation which is observed in a subset of ALS cases and has been shown to directly correlate with TDP-43 pathology in some brain regions. Additionally, TDP-43 proteinopathy is observed outside of ALS which may complicate the interpretation of case vs control approaches to target discovery. Here, we propose a pathology-stratified approach to empower targeted theranostics. We hypothesised that biologically distinct ALS subtypes may be defined by specific metabolic dysfunction linked to brain-accumulated ferritin and TDP-43 pathology.

**Methods:** Post-mortem primary motor cortex tissue from 15 ALS cases and 20 age- and sex-matched controls was stratified, using immunohistochemistry, by single- or co-occurrence of ferritin accumulation, and pathological TDP-43. Untargeted metabolomics (>1,000 metabolites) was performed, and samples were stratified into dual positive (ferritin and TDP-43), single positive (either), or negative. Group-discriminating metabolites were identified using partial least squares discriminant analysis.

**Results:** Dual ferritin and TDP-43 pathology reflected a distinct metabolomic profile, separable from single-pathology states. This dual positive metabolic signature was characterised by disruption of lysophospholipid, lysoplasmalogen, and fatty acid metabolism, consistent with impaired membrane and energy homeostasis. In contrast, pathological TDP-43 presence without ferritin, was characterised metabolically by significant depletion of secondary bile acids and increase in glycosylation markers, whilst ferritin accumulation alone reflected significant increase in oxidative stress and depletion of lipid peroxidation inhibition markers. The dual positive state suggests failure of compensatory metabolic responses present in single-pathology conditions.

**Conclusions:** Ferritin accumulation and TDP-43 pathology define biologically distinct subtypes associated with ALS with divergent metabolic vulnerabilities. The metabolic signature associated with dual pathology provides a mechanistic correlate to MRI-visible ferritin accumulated iron, supporting paired non-invasive biomarker and target discovery for pathology-dependent patient stratification. These findings argue for pathway-targeted, subtype-specific therapeutic strategies and highlight the necessity of precision medicine approaches in ALS.

**Short abstract:** Amyotrophic lateral sclerosis (ALS) exhibits profound molecular heterogeneity that is not captured by current clinical classifications. Additionally, TDP-43 proteinopathy is observed outside of ALS which may complicate the interpretation of case vs control approaches to target discovery. Here, we propose a pathology-stratified approach to therapeutic target discovery, identifying convergent iron dysregulation and TDP-43 pathology with specific metabolic consequences. Post-mortem primary motor cortex tissue from 15 ALS cases and 20 controls was investigated for ferritin, and pathological TDP-43 using RNA aptamer-based immunostaining. Untargeted metabolomics (>1,000 metabolites) was performed with stratification into dual positive, single positive, or negative groups, followed by partial least squares discriminant analysis. Dual ferritin and TDP-43 pathology produced a distinct metabolic state characterised by disruption of lysophospholipid, lysoplasmalogen, and fatty acid metabolism, indicating impaired membrane integrity and energy homeostasis. In contrast, single positive states engaged divergent compensatory pathways involving bile acid metabolism, glycosylation, or oxidative stress regulation. Ferritin–TDP-43 convergence defines a metabolically decompensated ALS subtype corresponding to MRI signatures, providing a mechanistic basis for imaging-guided, pathology-dependent patient stratification and targeted intervention.

**Key Findings:**

- Metabolically distinct subtypes were defined by the presence or absence of ferritin-associated iron accumulation and TDP-43 pathology in the primary motor cortex.
- Concurrent ferritin and TDP-43 pathology produce a unique, metabolically decompensated state characterised by disrupted lipid, membrane, and energy metabolism, distinct from either pathology alone.
- Single positive states engage divergent compensatory metabolic pathways, which are lost when ferritin and TDP-43 co-occur.
- The metabolic signature of dual positivity provides a mechanistic correlate to the MRI-visible motor band sign.
- These findings support the use of pathology-based stratification of ALS patients and a foundation for pathway-targeted, precision therapeutic approaches.

**Graphical Abstract:** 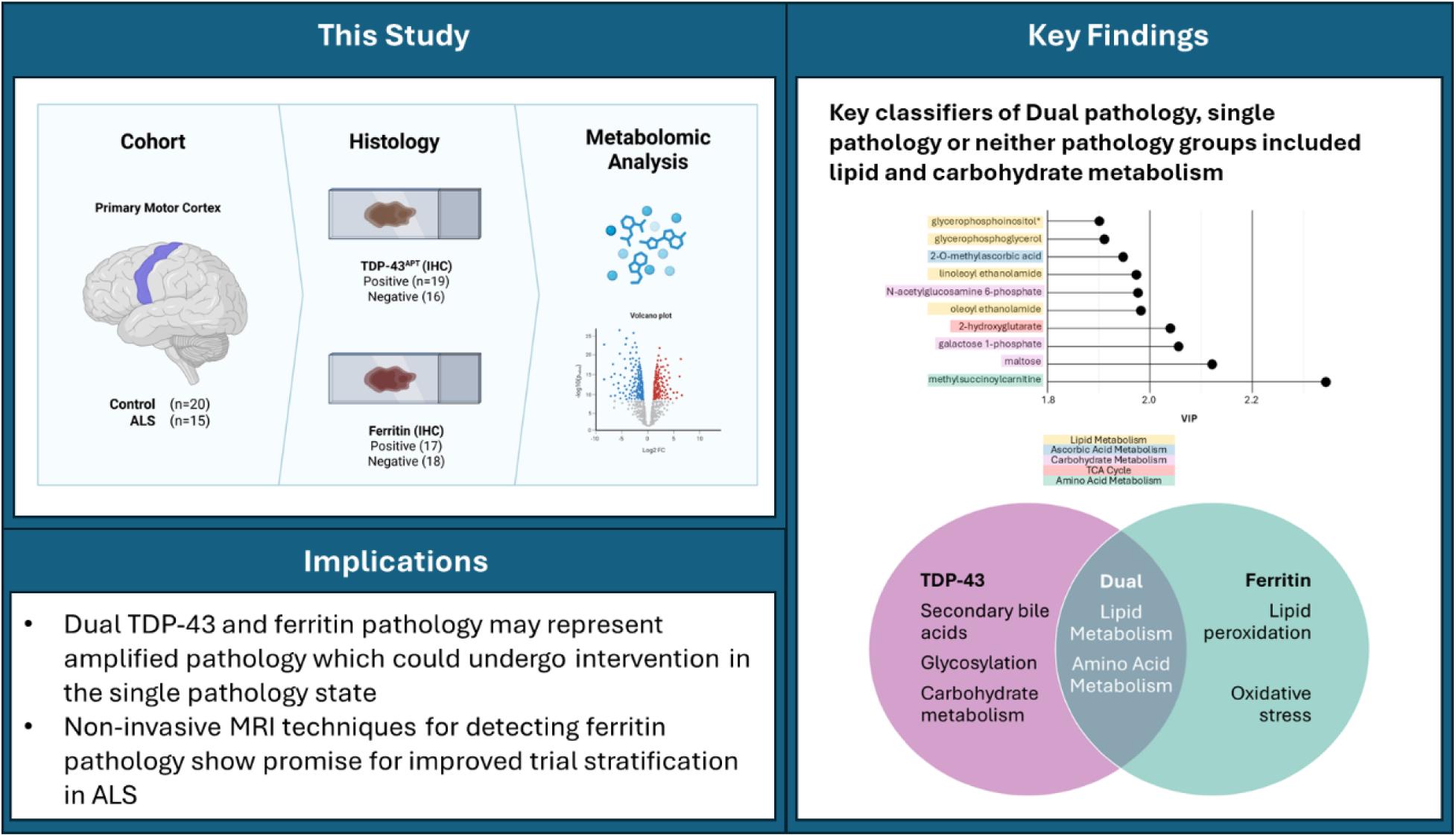

## Introduction

Amyotrophic lateral sclerosis (ALS) is recognised as a clinically and biologically heterogeneous disease. Indeed, recent large-scale molecular profiling studies have identified multiple distinct ALS endophenotypes, underscoring that convergent clinical features likely arise from divergent underlying biological processes [1–3]. This heterogeneity presents a major obstacle to biomarker development and therapeutic efficacy, as disease-modifying interventions are unlikely to be uniformly effective across molecularly distinct subgroups. Accordingly, a central challenge in ALS research is the development of stratification strategies that capture biologically meaningful variation and enable targeted therapeutic approaches. Achieving this goal requires biomarkers that are both mechanistically informative and amenable to clinical implementation.

Pathological transactive response DNA-binding protein 43 (TDP-43) aggregation represents a core molecular feature of ALS, yet its presence is not restricted to this disease. TDP-43 pathology is observed across a spectrum of neurodegenerative conditions, including Alzheimer’s disease, and is also detected in a subset of cognitively normal older individuals [4–7]. These observations challenge traditional disease-centric frameworks and motivate a shift toward pathology-driven stratification. Recent methodological advances, including the development of a highly sensitive TDP-43 RNA aptamer and probes and antibodies detecting cryptic splicing events, now permit robust, disease-agnostic detection of pathological TDP-43 [8–9]. These tools provide an opportunity to redefine TDP-43 pathology as a stratifying biomarker rather than a binary diagnostic feature.

Region-specific accumulation of ferritin-bound iron has also emerged as a promising biomarker in ALS, particularly due to its detectability using non-invasive MRI [10–11]. In the primary motor cortex, this pathology manifests as MRI hypointensity, the so-called *motor band sign* (MBS), and shows disease specificity for ALS [11]. Ferritin accumulation is a histological correlate of motor band sign and has exhibited region-dependent associations with TDP-43 pathology [4,12]. Importantly, however, ferritin accumulation and motor band sign are observed in only a subset of ALS cases [4,11,13], suggesting detection of a distinct molecular subtype rather than a universal feature of disease. This raises the possibility that ferritin accumulation identifies a biologically coherent subgroup of patients with shared downstream disease mechanisms and therapeutic vulnerabilities.

To date, metabolomic studies in ALS have largely adopted a *case vs control* framework, identifying alterations in energy and lipid metabolism in people with ALS relative to healthy age-and-sex matched controls [14–18]. However, such approaches may obscure biologically meaningful signals arising from molecular heterogeneity, particularly given the presence of TDP-43 pathology beyond classical disease boundaries [4–5,8–9]. Here, we adopt a precision, pathology-informed approach to interrogate the metabolic consequences of brain ferritin and TDP-43 pathology. Specifically, we assess metabolomic profiles of post-mortem primary motor cortex tissue from ALS cases and non-neurological controls stratified by the presence or absence of ferritin accumulation and pathological TDP-43. By defining the metabolic signatures associated with these pathological features, we aim to establish whether MRI-detectable ferritin accumulation serves as a mechanistically informative, non-invasive biomarker of a distinct ALS endophenotype. This work seeks to bridge imaging, molecular pathology, and metabolism to advance biologically grounded patient stratification in ALS.

## Methods

### Study cohort and design

A cohort of post-mortem tissue from the primary motor cortex of 35 individuals consisting of ALS cases (n=15; determined by pathological diagnosis) and non-neurological disease controls (n=20) was obtained from the National Institute for Health (NIH) NeuroBioBanks at the Mount Sinai NIH Brain and Tissue Repository (NBTR); the University of Maryland Brain and Tissue Bank, Baltimore, MD; Harvard Brain Tissue Resource Center (HBTRC) and the University of Miami Brain Endowment Bank^TM^ (Table 1) . Formalin fixed paraffin embedded (FFPE) tissue and paired frozen tissue was obtained. NIH NeuroBioBanks are operated under their respective institution’s internal review board approvals and obtain informed consent for brain donation. Our study focuses on stratification by molecular pathology rather than a disease vs control approach in order to determine biologically meaningful signals that may be obscured by molecular heterogeneity in ALS. To do this, we focus on two key molecular pathologies, TDP-43 loss- and gain-of-function detected by a TDP-43 RNA aptamer and ferritin accumulation indicative of motor band sign, detected by a ferritin antibody. To account for known variation in presence of each of these molecular pathologies in the “healthy control” population [4–7] a large cohort of 20 non-neurological disease controls were included in this study from multiple biobanks (Table 1). Partial least squares discriminant analysis (PLS-DA) was implemented to identify the metabolites most important for classification of pathology groups.

**Table 1.** Study demographics. Cases were age- and sex-matched by disease status with no significant difference in age (p=0.867; Wilcox test) and sex (p=0.522; χ^2^ test) between those with pathological diagnosis of ALS and controls. *ALS = Amyotrophic Lateral Sclerosis; AD = Alzheimer’s Disease; F = Female; M – Male; T+F+ = TDP-43^APT^ pathology positive, Ferritin high; T+F- = TDP-43^APT^ pathology positive, Ferritin low; T-F+ = TDP-43^APT^ pathology negative, Ferritin high; TF- = TDP-43^APT^ pathology negative, Ferritin low*.

| Case Number | Disease Status | Brain Bank | Age | Sex | Race | PMI (Hours) | Pathology Group |
| --- | --- | --- | --- | --- | --- | --- | --- |
| 1 | ALS | Maryland | 74 | F | Caucasian | 19 | T+F- |
| 2 | ALS | Maryland | 75 | M | Caucasian | 27 | T+F+ |
| 3 | ALS | Miami | 71 | F | Caucasian | 28 | T+F- |
| 4 | ALS | Miami | 81 | M | Caucasian | 32.42 | T+F- |
| 5 | ALS | Miami | 74 | M | Caucasian | 20.2 | T+F- |
| 6 | ALS | Miami | 64 | F | Caucasian | 47.5 | T+F- |
| 7 | ALS | Miami | 80 | M | Caucasian | 9.3 | T+F- |
| 8 | ALS | Harvard | 77 | F | Caucasian | 22.33 | T+F- |
| 9 | ALS | Harvard | 54 | M | Caucasian | 24.17 | T+F- |
| 10 | ALS | Harvard | 70 | M | Caucasian | 33.87 | T+F- |
| 11 | ALS and AD | Mount Sinai | 76 | F | Caucasian | 20.67 | T-F+ |
| 12 | ALS and AD | Mount Sinai | 53 | M | Caucasian | 4.75 | T+F+ |
| 13 | ALS and AD | Harvard | 75 | M | Caucasian | 20.06 | T-F- |
| 14 | ALS and PD | Miami | 70 | M | Caucasian | 18.48 | T-F- |
| 15 | ALS and PD | Miami | 79 | M | Hispanic or Latino | 20.41 | T-F- |
| 16 | Control | Mount Sinai | 76 | F | Hispanic | 4.25 | T+F+ |
| 17 | Control | Mount Sinai | 79 | M | Caucasian | 16 | T-F- |
| 18 | Control | Mount Sinai | 75 | M | Caucasian | 16.63 | T-F+ |
| 19 | Control | Mount Sinai | 76 | M | Caucasian | 2.92 | T-F+ |
| 20 | Control | Mount Sinai | 67 | F | Caucasian | 16.5 | T-F+ |
| 21 | Control | Mount Sinai | 64 | F | Caucasian | 19.03 | T-F- |
| 22 | Control | Mount Sinai | 75 | F | Caucasian | 9.92 | T+F+ |
| 23 | Control | Mount Sinai | 81 | M | Caucasian | 11.67 | T-F- |
| 24 | Control | Mount Sinai | 76 | F | Caucasian | 9.33 | T-F- |
| 25 | Control | Miami | 58 | M | Caucasian | 20.6 | T+F- |
| 26 | Control | Miami | 53 | M | Hispanic or Latino | 17.6 | T-F+ |
| 27 | Control | Miami | 70 | M | Hispanic or Latino | 12.73 | T+F+ |
| 28 | Control | Miami | 59 | M | Caucasian | 25.5 | T+F+ |
| 29 | Control | Harvard | 78 | F | Caucasian | 35.17 | T-F+ |
| 30 | Control | Harvard | 80 | M | Caucasian | 15.5 | T-F+ |
| 31 | Control | Harvard | 78 | F | Caucasian | 23.38 | T-F+ |
| 32 | Control | Harvard | 74 | M | Caucasian | 26 | T+F+ |
| 33 | Control | Harvard | 72 | F | Unknown | 18.25 | T+F+ |
| 34 | Control | Harvard | 76 | F | Unknown | 8.08 | T-F+ |
| 35 | Control | Harvard | 70 | F | Caucasian | 21.33 | T+F- |

### Immunostaining

TDP-43 aptamer [8] staining was performed on FFPE tissue as is previously published [19]. TDP-43 aptamer staining was graded manually for presence and absence of pathology by a trained histopathologist (JMG) blinded to clinical information.

Immunostaining of ferritin was performed on FFPE tissue using the Novolink Polymer detection system with Abcam Recombinant anti-Ferritin antibody (ab287968) at 1 in 100 dilution with DAB chromogen and pretreatment for 10 minutes in Tris/EDTA buffer in a pressure cooker. Counterstaining was performed with haematoxylin, according to standard operating procedures.

Ferritin superpixel analysis was assessed using the freely available QuPath software implementing superpixel analysis using the following code:

selectAnnotations();

runPlugin(’qupath.imagej.superpixels.DoGSuperpixelsPlugin’, ’{"downsampleFactor": 1.0, "sigmaMicrons": 3, "minThreshold": 10.0, "maxThreshold": 230.0, "noiseThreshold": 0.0}’);

selectDetections(); runPlugin(’qupath.lib.algorithms.IntensityFeaturesPlugin’, ’{"downsample": 1.0, "region": "ROI", "tileSizePixels": 200.0, "colorOD": false, "colorStain1": false, "colorStain2": true, "colorStain3": false, "colorRed": false, "colorGreen": false, "colorBlue": false, "colorHue": false, "colorSaturation": false, "colorBrightness": false, "doMean": true, "doStdDev": false, "doMinMax": false, "doMedian": false, "doHaralick": false, "haralickDistance": 1, "haralickBins": 32}’);

setDetectionIntensityClassifications("ROI: 2.00 µm per pixel: DAB: Mean", 0.3602, 0.4091, 0.4580)

Number of 1+ detections (Mean+SD), more intense 2+ detections (Mean+2SD) and most severe 3+ detections (Mean+3SD) were quantified and the weighted score (1x number of 1+ detections + 2x number of 2+ detections + 3x number of 3+ detections) was calculated.

### Stratification

ALS cases and controls were age and sex matched (Table 1). A disease agnostic, pathology focused approach was then taken to determine changes associated with single and concurrent pathology. Four pathology groups were initially identified based on absence and presence of TDP-43 pathology determined by a pathologist (JMG); (i) negative for both TDP-43 and ferritin pathology; (ii) Negative for TDP-43 pathology and positive for ferritin pathology; (iii) positive for TDP-43 pathology and negative for ferritin pathology, (iv) positive for both TDP-43 and ferritin pathology. Ferritin accumulation rated as high or low, determined by thresholding of superpixel quantitative analysis (Figure 1). Thresholding on a larger cohort of 60 primary motor cortex samples from cases with Alzheimer’s disease (n=10), Parkinson’s disease (n=10), ALS (n=10), multiple neurological diagnoses (n=10) and non-neurological controls (n=20) determined a cut-off for discriminating high and low ferritin at the median ferritin superpixel score across all 60 cases (787).

**Figure 1.**
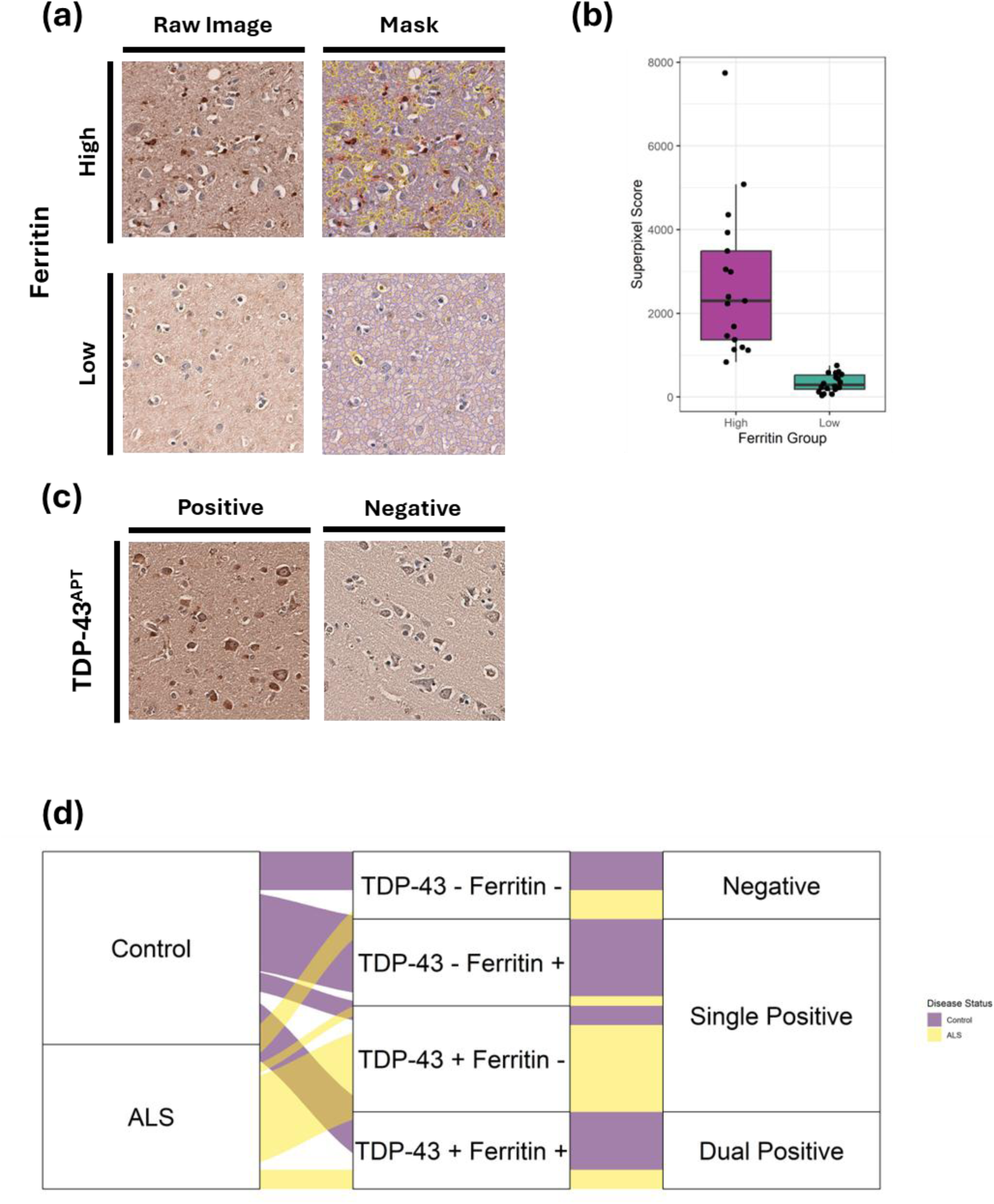
**(a)** Representative images show raw images and superpixel masks of ferritin staining in individuals with high and low ferritin staining intensity. Superpixel masks show blue tiles with intensity ≤ mean intensity across wider cohort, yellow tiles with intensity > 1 standard deviation above mean intensity across wider cohort and red tiles with intensity > 2 standard deviations above the mean intensity across wider cohort. Images acquired at 20X magnification on EVOS M500 microscope. **(b)** Superpixel score distribution of ferritin staining intensities by group. **(c)** Representative images show TDP-43^APT^ staining in individuals with presence and absence of TDP-43 pathology as rated by a pathologist. Images acquired at 20X magnification on EVOS M500 microscope. **(d)** Alluvial plot showing distribution of ALS and control individuals in each of the four pathology groups. ALS cases negative for TDP-43^APT^ pathology were all cases with multiproteinopathy where pathology reports describe TDP-43 pathology outside of the motor cortex.

### Metabolomics

An equal mass of each frozen tissue sample was analysed on Metabolon’s Global Discovery platform. Samples were prepared using the automated MicroLab STAR® system (Hamilton Company). Several recovery standards were added prior to the first step in the extraction process for QC purposes. To remove protein, dissociate small molecules bound to protein or trapped in the precipitated protein matrix, and to recover chemically diverse metabolites, proteins were precipitated with methanol under vigorous shaking for 2 min (Glen Mills GenoGrinder 2000) followed by centrifugation. The resulting extract was divided into multiple fractions: two for analysis by two separate reverse phase (RP)/UPLC-MS/MS methods with positive ion mode electrospray ionization (ESI), one for analysis by RP/UPLC-MS/MS with negative ion mode ESI, one for analysis by HILIC/UPLC-MS/MS with negative ion mode ESI, while the remaining fractions were reserved for backup. Samples were placed briefly on a TurboVap® (Zymark) to remove the organic solvent. The sample extracts were stored overnight under nitrogen before preparation for analysis.

### Ultrahigh Performance Liquid Chromatography-Tandem Mass Spectroscopy (UPLC-MS/MS)

All described metabolite identification methods utilized a Waters ACQUITY ultra-performance liquid chromatography (UPLC) and a Thermo Scientific Q-Exactive high resolution/accurate mass spectrometer interfaced with a heated electrospray ionization (HESI-II) source and Orbitrap mass analyser operated at 35,000 mass resolution (PMID: 32445384). The dried sample extract was then reconstituted in solvents compatible to each of the four methods hereafter described. Each reconstitution solvent contained a series of standards at fixed concentrations to ensure injection and chromatographic consistency. One aliquot was analysed using acidic positive ion conditions, chromatographically optimized for more hydrophilic compounds (PosEarly). In this method, the extract was gradient eluted from a C18 column (Waters UPLC BEH C18-2.1x100 mm, 1.7 µm) using water and methanol, containing 0.05% perfluoropentanoic acid (PFPA) and 0.1% formic acid (FA). Another, second aliquot was also analysed using acidic positive ion conditions, however it was chromatographically optimized for more hydrophobic compounds (PosLate). Here, the extract was gradient eluted from the same C18 column using methanol, acetonitrile, water, 0.05% PFPA and 0.01% FA and was operated at an overall higher organic content. A third aliquot was analysed using basic negative ion optimized conditions using a separate dedicated C18 column (Neg). The basic extracts were gradient eluted from the column using methanol and water, however with 6.5mM Ammonium Bicarbonate at pH 8. The fourth aliquot was analysed via negative ionization following elution from a HILIC column (Waters UPLC BEH Amide 2.1x150 mm, 1.7 µm) using a gradient consisting of water and acetonitrile with 10mM Ammonium Formate, pH 10.8 (HILIC). The MS analysis alternated between MS and data-dependent MSn scans using dynamic exclusion. The scan range varied slightly between methods but covered 70-1000 m/z.

### Statistical analysis

For metabolites with values below the detection limit, missing values were imputed using the minimum observed value for each compound. Log transformation was used to normalise metabolite levels. Welch’s two-sample t-tests were used to identify significant differences in biochemicals between experimental groups. An estimate of the false discovery rate (q-value) was calculated to take into account multiple comparisons. Volcano plots and VIP plots were made using ggplot2 packages in R [20].

Metabolites where all samples had imputed value were excluded from the data matrix. Supervised partial least squares discriminant analysis (PLS-DA) was performed and validated on the resulting data matrix split into high and low ferritin groups using the R package mixOmics [21]. Validation of the optimal number of components was performed based on a distance matrix approach. Repeated cross-validation (10 × 3−fold CV) was used to evaluate the PLS-DA classification performance (OER and BER), for each type of prediction distance; maximum distance, centroid distance and Mahalanobis distance (Supplementary Fig. 1). Metabolite loading on each component was assessed to determine pathways important for discriminating high and low ferritin groups. Variable importance in projection (VIP) score analysis on the PLS-DA further validated classifying metabolites where a stringent VIP>2 identified top classifying metabolites. Area under the curve (AUC) and receiver operating characteristics (ROC) were used to further validate the model.

## Results

### Stratification by ferritin and TDP-43 pathological burden defines distinct cohort subgroups

Stratification based on ferritin accumulation, indicative of motor band sign presence, identified 17 individuals with ferritin burden above the discriminating threshold and 18 individuals with ferritin burden below the discriminating threshold (Figure 1A, B & D). Absence vs presence rating of TDP-43 pathology indicated 19 individuals had TDP-43 pathology and 16 individuals did not (Figure 1C & D). An important note is that all cases with pure ALS diagnosis had TDP-43 pathology, whilst those with ALS combined with either Alzheimer’s disease or Parkinson’s disease tended to be negative for TDP-43 pathology in the BA4 region assessed (Table1). However, where pathology reports for these individuals were available, pathological ALS diagnosis was reached due to TDP-43 pathology outside of the BA4 region.

### PLS-DA model classifies absence or presence of TDP-43 and ferritin pathology with strong model performance

To identify key discriminating metabolites between TDP-43 and ferritin pathology groups, a partial least squares discriminant analysis (PLS-DA) was adopted. The optimal number of components (2) was determined by calculating the classification error (OER and BER) for models with up to 10 components (Supplementary figure 1). An initial four group PLS-DA model demonstrated a lack of distinct metabolomic profile between those with a single pathology of either TDP-43 or ferritin pathology (Figure 2A). Therefore, a three-group model was adopted: (i) Negative (neither TDP-43 or ferritin pathology), (ii) Single positive (presence of either TDP-43 or ferritin pathology) and (iii) Dual positive (presence of both TDP-43 and ferritin pathology). There was no significant difference in age (p>0.05; Kruskal-Wallis; Figure 2B), disease status (p>0.05; χ^2^ test; Figure 2C), sex (p>0.05; χ^2^ test; Figure 2D) or brain bank status between groups (p>0.05; χ^2^ test; Figure 2E).

**Figure 2.**
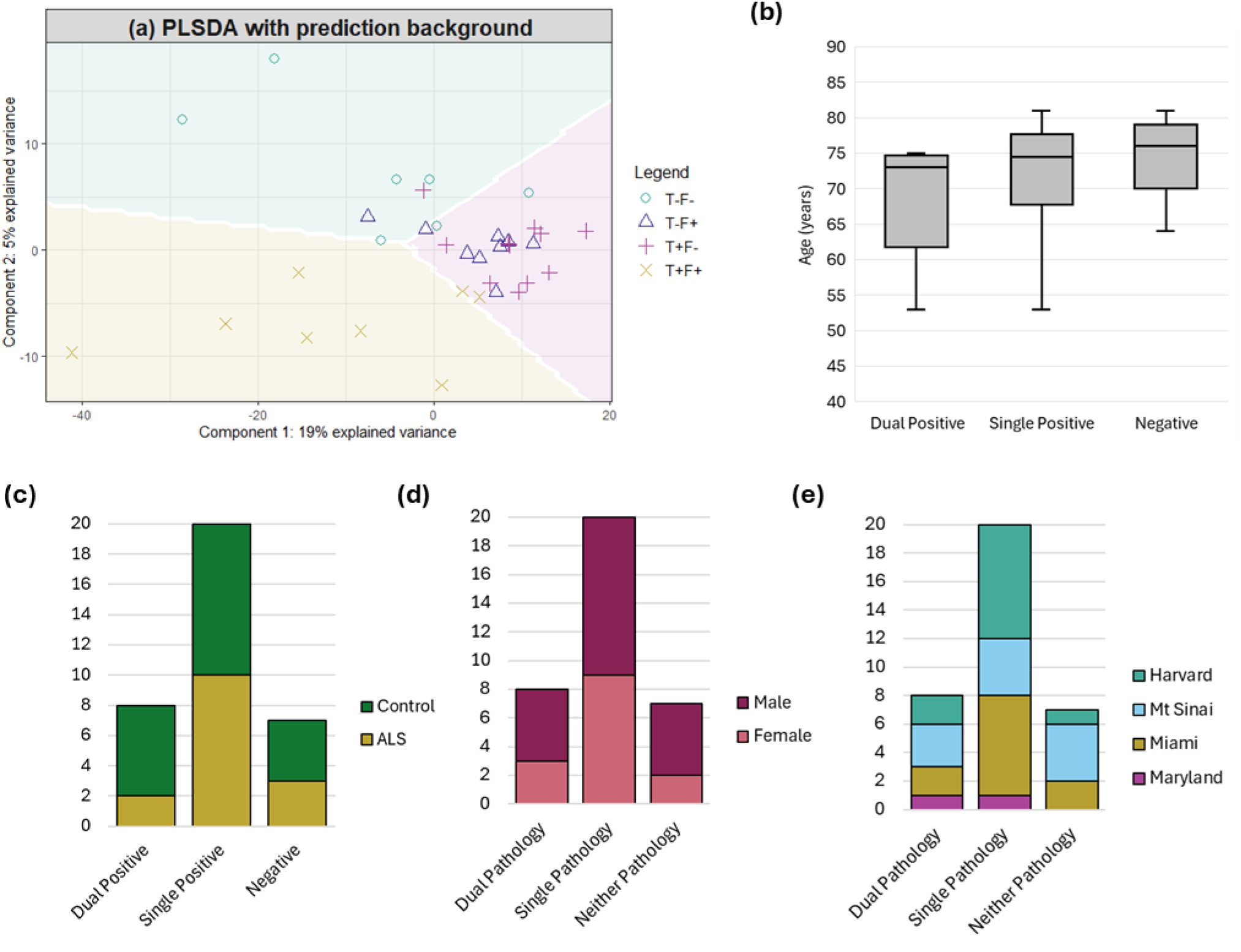
**(a)** Two component PLS-DA model reveals metabolomic discrimination between those with both TDP-43 and ferritin pathology (dual positive), either TDP-43 or ferritin pathology (single positive) and no TDP-43 or ferritin pathology (negative). Individuals with single positivity (either TDP-43 or ferritin in the BA4 region form an indistinct metabolomic profile. When data was instead split into three groups (i) dual positive; (ii) single positive; (iii) negative, there was no significant difference in **(b)** age, **(c)** disease status, **(d)** sex or **(e)** brain bank status between the groups.

A PLS-DA classifying three pathology groups demonstrated significant discrimination between groups with combined explained variance of 25% (Figure 3A & B). Component 1 demonstrates significant model sensitivity and specificity for discrimination of dual positive and single positive from other groups (AUC=0.82, p<0.01 & AUC=0.88, p<0.001 respectively; Figure 3C), but not for discrimination of negative from other groups (AUC=0.73, p>0.05; Figure 3C). When combined, the two-component model demonstrates significant specificity and sensitivity in classification performance across all groups (AUC>0.89, p<0.001; Figure 3D). Component 2 therefore represents metabolites important for classifying negative individuals, whilst component 1 represents metabolites important for classifying dual positive from single positive individuals.

**Figure 3.**
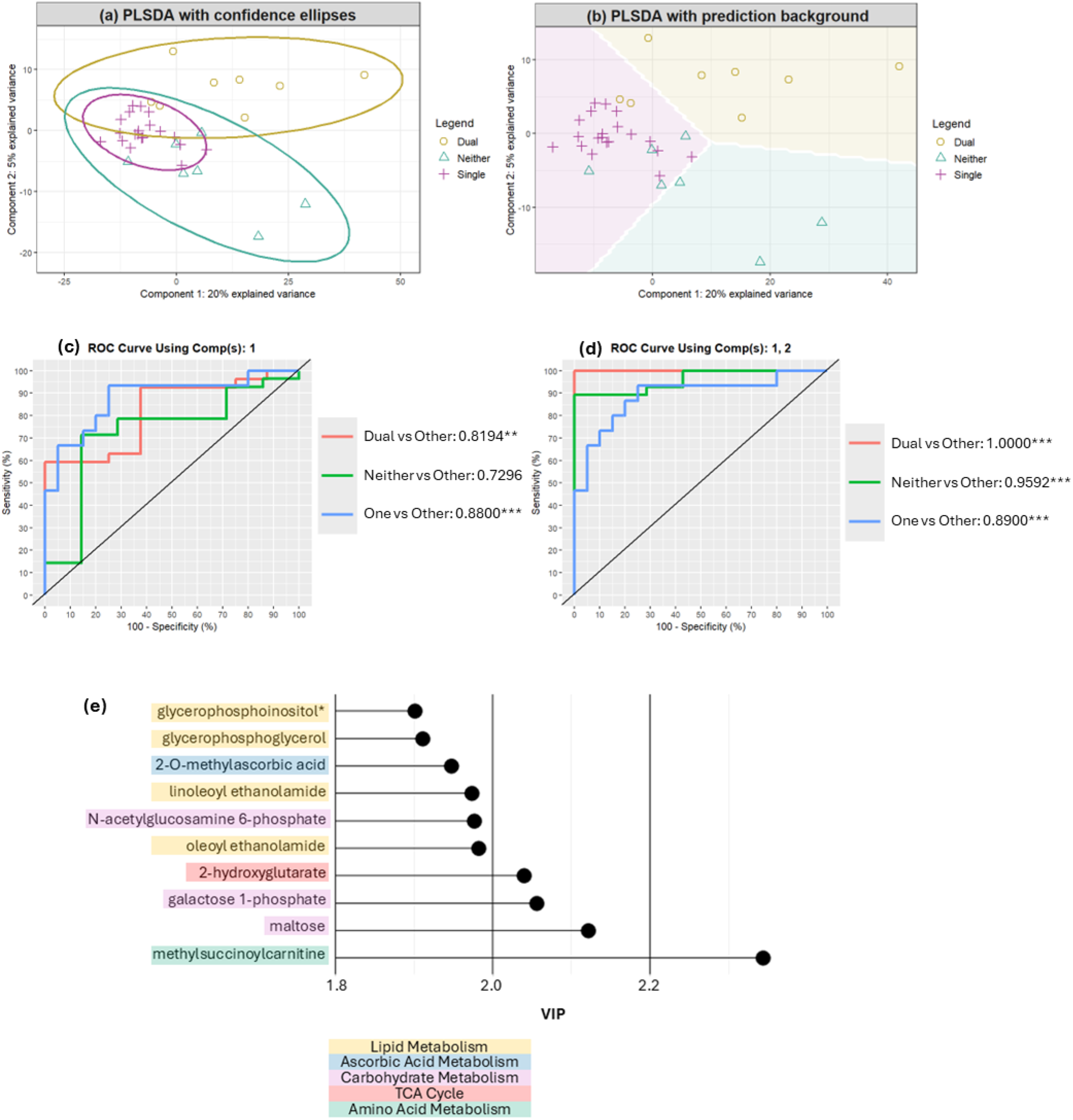
(a-b) Sample plots of metabolomics data after a 3 group, 2 component PLS-DA model was operated. **(a)** Sample separation using 1^st^ and 2^nd^ latent component scores with the confidence ellipses (95% CE) of each class. **(b)** Sample separation using 1^st^ and 2^nd^ latent component scores with prediction background generated by these samples. **(c-d)** ROC curve and AUC outcome from 3 group, 2 component PLS-DA. **(c)** Component 1 demonstrates significant discrimination between those with both pathologies and one pathology (AUC>0.8 and p<0.01). **(d)** Combined with component two, significant discrimination is reached between all groups (AUC >0.89, p<0.001). **(e)** Variable importance in projection (VIP) plot for metabolites with top 10 VIP scores. Colours represent metabolic pathway.

### Concurrent TDP-43 and ferritin pathology produce a distinct profile of disrupted lipid, membrane, and energy metabolism

Metabolites with top 10 variable importance in projection (VIP) scores associated with lipid metabolism indicate lower phospholipid levels but higher endocannabinoid levels in the negative group (Figure 3E; Supplementary figure 3). Other metabolites with highest VIP scores indicate increased levels of carbohydrates associated with glycosylation and glycan processing in the dual positive group (Figure 3E; Supplementary figure 3). Additionally, metabolites associated with disrupted fatty acid metabolism (2-hydroxyglutarate and methylsuccinoylcarnitine) demonstrate increases in those dual or single positivity (Figure 3E; Supplementary figure 3).

Component 1 which performs best for distinguishing dual positivity from single positivity is loaded most heavily by lipid alterations including increased levels of lysophospholipids and lysoplasmalogens in those with dual positivity, whilst changes in cysteine-glutathione metabolism and B vitamin metabolism markers are higher in those with single positivity compared to dual positivity (Figure 4A).

**Figure 4.**
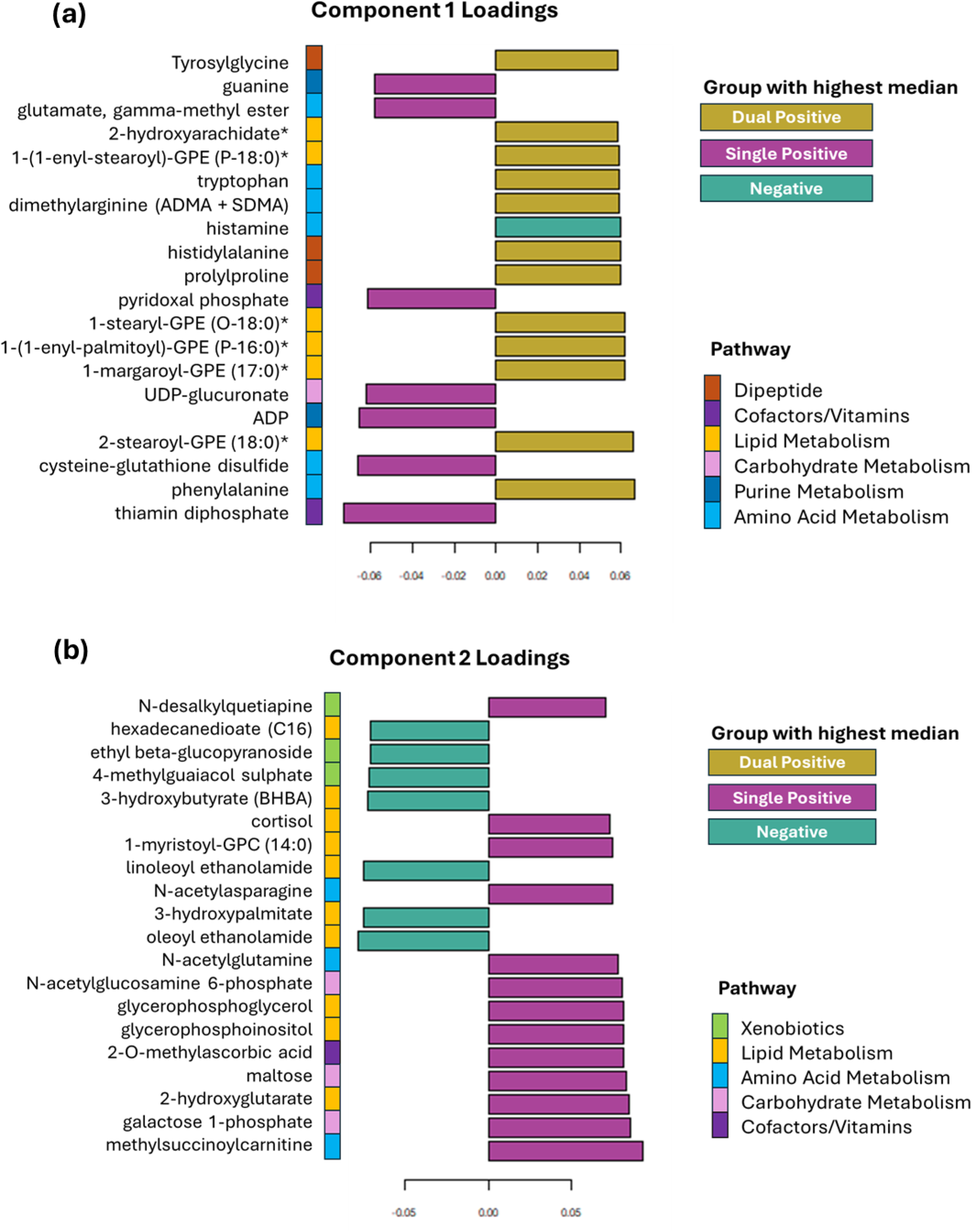
Principal component loadings plot with vector loadings of 20 highest loading variables **(a)** in component 1 and **(b)** component 2 of PLS-DA. Key contributing pathways include amino acid and lipid metabolism. Bar colour represents the group with the highest median level of each metabolite.

Component 2 which performs best for distinguishing negative subjects from those with a pathology is also loaded most heavily by lipid levels, however, here metabolites are indicative of impaired fatty acid and phospholipid metabolism (Figure 4B). Those negative for pathology exhibited higher levels of fatty acids, phospholipids and endocannabinoids, whilst dual positive individuals exhibited higher levels of metabolites associated with carbohydrate metabolism (Figure 4B).

### Single positive states engage divergent compensatory metabolic pathways, which are lost when ferritin and TDP-43 co-occur

We compared metabolites between those with and without ferritin pathology, separately to those with and without TDP-43 pathology to determine whether one specific pathology may be more strongly associated with single pathology metabolite changes observed in the PLS-DA model. Comparison between those with and without TDP-43 pathology demonstrates significantly altered carbohydrate metabolism and secondary bile acid metabolism changes (Figure 5A). Increases in galactose-1-phosphate and maltose and a decrease in galactonate associated with TDP-43 pathology presence were highly loaded metabolites in the PLS-DA discrimination model.

**Figure 5.**
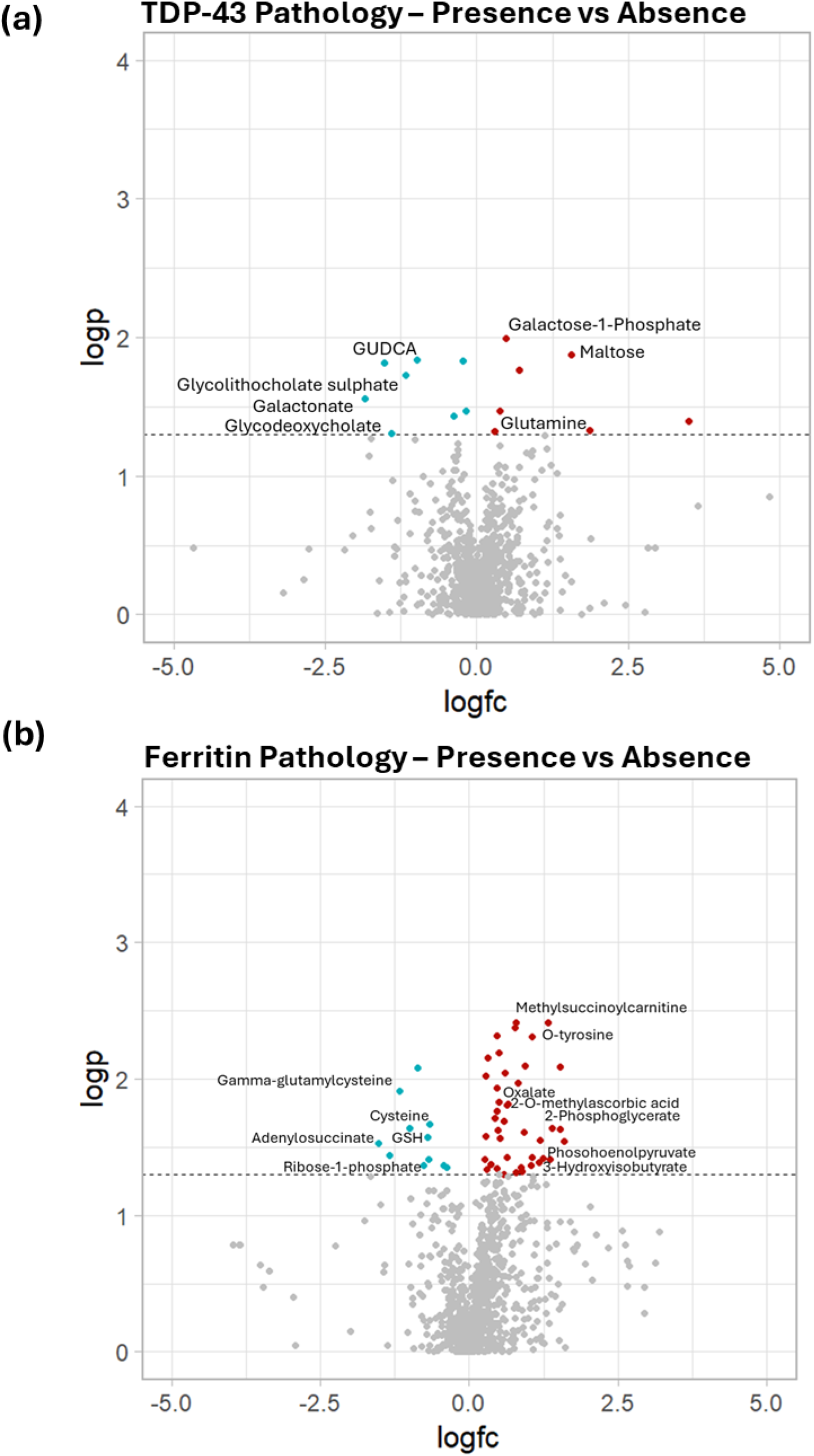
Volcano plots to show metabolites with significantly different levels between **(a)** TDP-43 positive vs TDP-43 negative labelled with metabolites related to carbohydrate and bile acid metabolism pathways and **(b)** Ferritin positive vs ferritin negative labelled with metabolites related to lipid peroxidation pathways.

Comparison of those with and without ferritin pathology exhibits alterations in several metabolites associated with impaired lipid peroxidation pathways, with lower levels of lipid peroxidation inhibiting metabolites (e.g. cysteine, glutathione, gamma-glutamylcysteine) and higher levels of reactive oxygen species (ROS) and lipid peroxidation (2-phosphoglycerate and oxalate). Metabolites associated with lipid alterations were also highly loaded in the PLS-DA model.

As several metabolites associated with cysteine-glutathione metabolism, ROS and fatty acid metabolism were highly loaded in component 1 of the PLS-DA model associated with single positivity, we investigated glutathione peroxidase 4 (GPX4) as a potential linking and targetable enzyme involved in this pathway. We demonstrate significant negative associations between GPX4 and ferritin superpixel burden score in the primary motor cortex of the individuals in this cohort (p=0.005, R=-0.50 respectively; Figure 6A). TDP-43 and GPX4 exhibit a quadratic relationship whereby GPX4 is either low or high in those with high TDP-43 pathology (R^2^=0.2434, p=0.0088; Figure 6B). There were no significant linear associations between GPX4 and TDP-43 (Supplementary figure 4). This indicates impaired lipid peroxidation inhibition in those with high ferritin levels.

**Figure 6.**
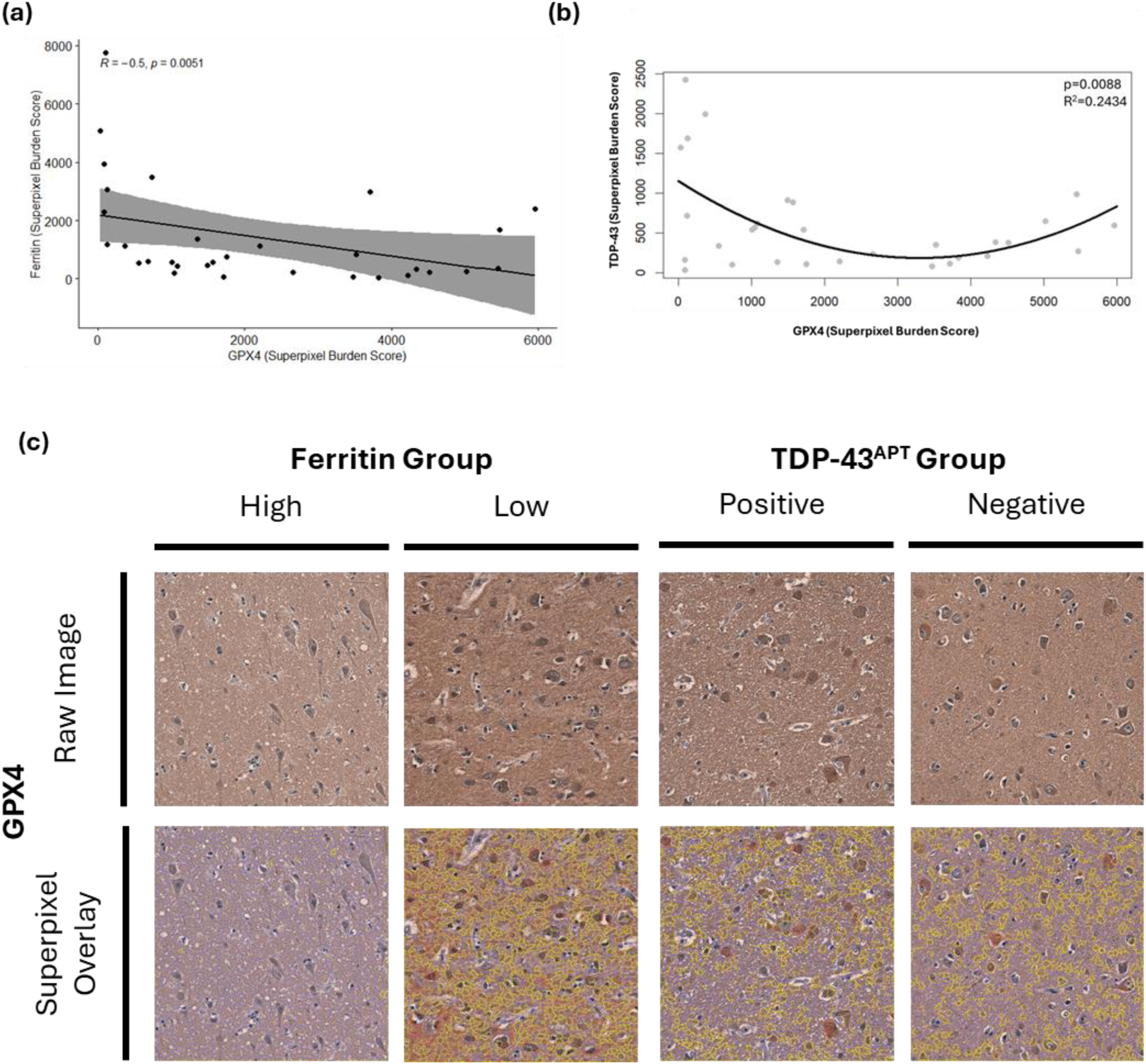
**(a)** Glutathione peroxidase 4 (GPX4) and ferritin in the BA4 region are significantly inversely correlated whereby individuals with higher ferritin had reduced levels of GPX4 when assessing superpixel burden score (Spearman; R=-0.5; p=0.0051). **(b)** GPX4 and TDP-43 in the BA4 region exhibit a significant quadratic relationship when assessing superpixel burden score (R^2^=0.24, p=0.0088) **(c)** Representative images show GPX4 staining across pathology groups. Superpixel masks show blue tiles with intensity ≤ mean intensity across wider cohort, yellow tiles with intensity > 1 standard deviation above mean intensity across wider cohort and red tiles with intensity > 2 standard deviations above the mean intensity across wider cohort. Raw images acquired at 20X magnification on EVOS M500 microscope.

## Discussion

ALS is increasingly recognised as a biologically heterogeneous disease in which multiple pathological and molecular mechanisms converge to produce a similar clinical phenotype. In this study, we applied a disease-agnostic, pathology-focused stratification strategy to examine metabolomic changes associated with ferritin accumulation and TDP-43 pathology in the primary motor cortex. Previous metabolomic studies in ALS have assumed that controls have no underlying pathology and that ALS cases are all the same, stratification based on TDP-43 and ferritin pathology presence revealed patterns that would not be evident under these previous conventional case–control approaches that assume biological homogeneity within ALS. Three key findings emerge from this work: (1) dual ferritin and TDP-43 pathology defines a distinct metabolically decompensated state, (2) single pathology states reveal distinct mechanistic pathways of compensation, and (3) ferritin accumulation provides a metabolic–imaging bridge for precision medicine in ALS.

### Dual ferritin and TDP-43 pathology defines a distinct metabolically decompensated state

We demonstrate that the presence of either TDP-43 pathology or ferritin accumulation alone is associated with relatively indistinct metabolomic profiles. In contrast, individuals with concurrent ferritin and TDP-43 pathology exhibit a distinct metabolomic signature, characterised by coordinated alterations in lipid, amino acid, and carbohydrate metabolism consistent with disruption of signalling and energy-producing pathways. This suggests that ferritin and TDP-43 pathologies may arise through partially independent mechanisms, but that their co-occurrence results in convergent metabolic failure that overwhelms cellular compensatory pathways.

Consistent with this interpretation, the dual-pathology group exhibited marked changes in lysophospholipids and lysoplasmalogens, with glycerophosphoinositol and glycerophosphoglycerol among the highest variable-importance (VIP) scoring metabolites. These phospholipids were elevated in individuals with pathology compared with those without pathology, consistent with previous metabolomic studies showing prominent lysophospholipid alterations in ALS [15,17]. Along with carnitines, lysophospholipids and genes involved in lipid and iron metabolism have previously been associated with disease progression and severity in ALS [17,22]. In the present study, these markers were increased in pathology-positive individuals, with the largest elevations occurring in those with dual pathology, suggesting that this state may represent a later or more metabolically compromised stage of disease.

Fatty-acid amides including linoleoyl ethanolamide (LE) and oleoyl ethanolamide (OE) also showed high VIP scores and were comparatively elevated in individuals without pathology. These molecules have previously been shown to inhibit NF-κB signalling and exert anti-inflammatory effects [23], raising the possibility that their depletion in pathology-positive individuals reflects loss of endogenous neuroprotective mechanisms. LE is also a precursor to linoleic acid, a fatty acid reported to be depleted in ALS, supplementation of which recently improved survival in a C9orf72 Drosophila model [14]. Together, these findings suggest that lipid dysregulation and impaired anti-inflammatory signalling represent key components of the metabolically decompensated dual-pathology state.

### Single pathology states reveal distinct mechanistic pathways

When ferritin and TDP-43 pathologies were examined independently, divergent metabolic signatures emerged, suggesting that each pathology engages different compensatory pathways prior to the metabolic collapse observed in the dual-positive state. TDP-43 pathology was primarily associated with alterations in carbohydrate and purine metabolism. Increased maltose, galactose-1-phosphate and N-acetylglucosamine-6-phosphate were observed in individuals with TDP-43 pathology, consistent with impaired energy-producing and glycosylation pathways. Increased maltose has previously been associated with ALS presence and progression [24], whilst N-acetylglucosamine-6-phosphate has been implicated in disrupted O-GlcNAcylation pathways across neurodegenerative diseases [25–26]. Secondary bile acid metabolism was also altered, consistent with previous observations that bile acids are depleted in ALS. Tauroursodeoxycholic acid (TUDCA) has progressed to phase III clinical trials for ALS (NCT03800524), although despite early promise [27], these trials ultimately failed to meet their primary endpoints, suggesting that simple replacement of depleted bile acids may be insufficient to address the underlying metabolic disruption.

Purine metabolism was also altered in TDP-43-positive individuals, particularly within guanine metabolism, with increased guanine and GDP levels observed. S-adenosylhomocysteine (SAH) was identified as a key contributing metabolite to this distinct metabolic profile and is indicative of DNA hypermethylation associated with TDP-43 proteinopathy [28]. SAH accumulation may also disrupt downstream purine metabolism and glycolytic regulation, further linking TDP-43 pathology with broader metabolic perturbations. In contrast, ferritin accumulation was specifically associated with markers of oxidative stress and mitochondrial dysfunction, including oxalate, methylsuccinylcarnitine, and O-tyrosine. These findings are consistent with the proposed oxidative stress molecular subtype of ALS (ox-ALS). Calcium signalling is considered a defining feature of this phenotype [29] and has been proposed to precede iron accumulation in individuals exhibiting the motor band sign [30–31]. Our findings therefore support the hypothesis that ferritin accumulation in the motor cortex reflects an oxidative stress–driven disease process.

Consistent with this, glutathione-related metabolism was altered in individuals with ferritin pathology. Glutathione peroxidase (GPX4) changes have been widely reported in ALS, particularly within oxidative stress–associated disease phenotypes [3]. In our cohort, GPX4 levels were significantly negatively associated with ferritin pathology but not with TDP-43 pathology. Interestingly, a quadratic relationship was observed between GPX4 and TDP-43 pathology, suggesting that GPX4 depletion may occur at intermediate levels of TDP-43 burden. This raises the possibility that oxidative stress driven by ferritin accumulation may compromise the ability of cells to compensate for TDP-43-related stress. Supporting this interpretation, recent work demonstrated rescue of TDP-43 cryptic exon inclusion following treatment with N-acetyl cysteine through amelioration of ROS-induced TDP-43 pathology [32].

Carnitine metabolism emerged as another key pathway linking oxidative stress and lipid metabolism. The highest VIP-scoring metabolite associated with ferritin accumulation was methylsuccinoylcarnitine, alongside several other carnitine-related metabolites including linoleoylcarnitine (C18:2), oleoylcarnitine (C18:1), and docosapentaenoylcarnitine (C22:5n6), reflecting disruption of mitochondrial fatty-acid metabolism. Trimethyl-5-aminovalerate metabolism, capable of inhibiting carnitine biosynthesis, was also an important predictor of pathology group [33]. Carnitine metabolism regulates multiple pathways relevant to ALS pathophysiology, including calcium homeostasis, mitochondrial β-oxidation, insulin sensitivity, branched-chain amino acid metabolism, and redox balance [33]. Notably, carnitine can enhance Nrf2-GPX4 antioxidant signalling, and acetyl-L-carnitine is currently under investigation in a phase II/III clinical trial for ALS (NCT06126315).

### Ferritin accumulation provides a metabolic–imaging bridge for precision medicine in ALS

A major implication of these findings is the translational link between the metabolic phenotype identified here and the MRI-visible motor band sign, which reflects iron accumulation in the motor cortex. Ferritin accumulation detected histologically corresponds to susceptibility changes detectable by susceptibility-weighted MRI [10], suggesting that the metabolic phenotype described here has a direct imaging correlate. Historically, the motor band sign has been considered a relatively poor diagnostic marker for ALS due to limited sensitivity and specificity when applied across all patients. However, the present findings suggest that this interpretation may overlook its potential value as a stratification biomarker rather than a universal diagnostic tool. In the context of precision medicine, the ability to identify a biologically defined subgroup of ALS patients non-invasively could be transformative.

Importantly, most metabolomic studies in ALS have relied on traditional case–control comparisons that assume biological homogeneity among patients and treat controls as metabolically normal baselines. Our pathology-first approach demonstrates that this assumption may obscure meaningful disease mechanisms. By stratifying individuals based on ferritin accumulation and TDP-43 pathology, we reveal metabolic phenotypes that are not apparent under conventional analytical frameworks. In particular, the identification of a metabolically decompensated state associated with dual pathology suggests that metabolic failure may represent a convergence point for multiple pathological processes.

These findings therefore support a model in which ferritin accumulation marks an oxidative stress–associated metabolic subtype of ALS that can be detected using MRI, enabling patient identification prior to or during early stages of metabolic decompensation. In turn, this provides a framework for targeted therapeutic strategies. For example, individuals with ferritin accumulation may represent the subgroup most likely to benefit from interventions aimed at restoring redox balance or lipid metabolism, such as N-acetyl cysteine or carnitine supplementation.

## Conclusion

Together, our findings establish dual TDP-43 and ferritin pathology as a biologically distinct state characterised by convergent metabolic decompensation, particularly affecting lipid, energy, and redox pathways. Crucially, ferritin accumulation provides a bridge between molecular pathology and neuroimaging, linking this metabolic phenotype to the MRI-visible motor band sign.

This work therefore reframes the motor band sign not as a universal diagnostic marker but as a precision stratification tool capable of identifying a biologically defined subset of ALS patients. By integrating molecular pathology, metabolomics, and imaging biomarkers, this approach provides a framework for resolving ALS heterogeneity and for designing mechanism-informed therapeutic trials targeting specific disease pathways.

## Author Contributions

JMG conceptualized, secured primary funding for, and led the study with key input from HS. JMG and HS prepared the tissue request and coordinated sample logistics. Immunohistochemistry was performed by HS, FLR, and JMG. HS undertook digital image acquisition and curation. Data curation was performed by HS. Data analysis was performed by HS with input from FMW and JMG. HS drafted the initial manuscript; subsequent drafts were prepared by all authors and finalized by HS, FMW and JMG.

## Conflicts of Interest

The authors declare no conflicts of interest.

## Data sharing

All data are available in the manuscript, tables and supplementary data.

## Funding

The research for this manuscript has been supported by: (i) a Target ALS Early-Stage Clinician Scientist award to JMG (FS-2023-ESC-S2), (ii) an NHS Grampian grant to FMW (GCA25107); (iii) an NIH grant to JMG and employing HS and FR (R01NS127186). Funders had no role in study design, data collection, data analyses, interpretation, or writing the manuscript.

## Acknowledgements

The authors would like to thank Metabolon, the University of Aberdeen Microscopy and Histology Core Facility in the Institute of Medical Sciences, the NIH NeuroBioBanks and the patients and families of the donors without whom this study would not be possible.

## Supplementary Materials

**Supplementary figure 1.**
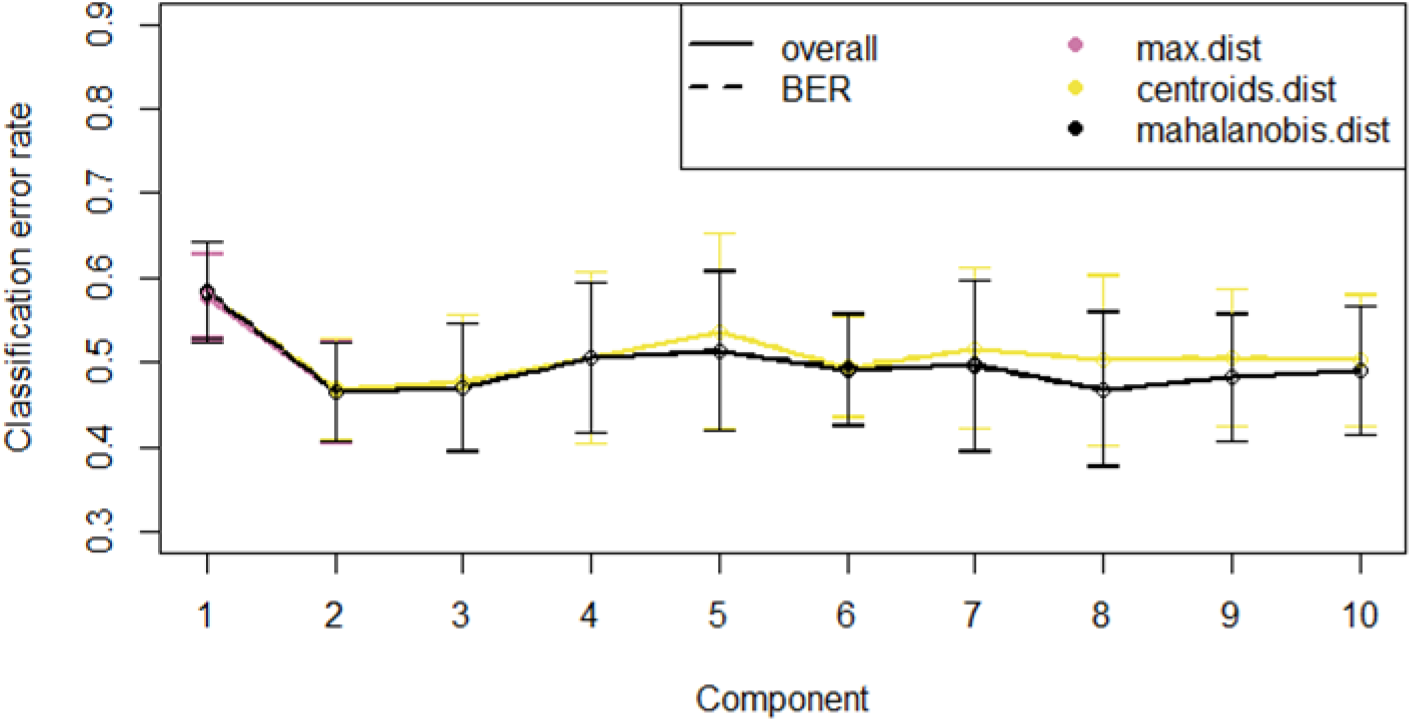
Tuning the number of components in PLS-DA demonstrates that for all prediction distances assessed, a 2-component model was optimal. For each component, repeated cross-validation (10 x 3-fold CV} was used to evaluate the PLS-DA classification performance (OER and BER), for each type of prediction distance; ’max.dist’, ’centroids.dist’ and ’mahalanobis.dist’.

**Supplementary figure 2.**
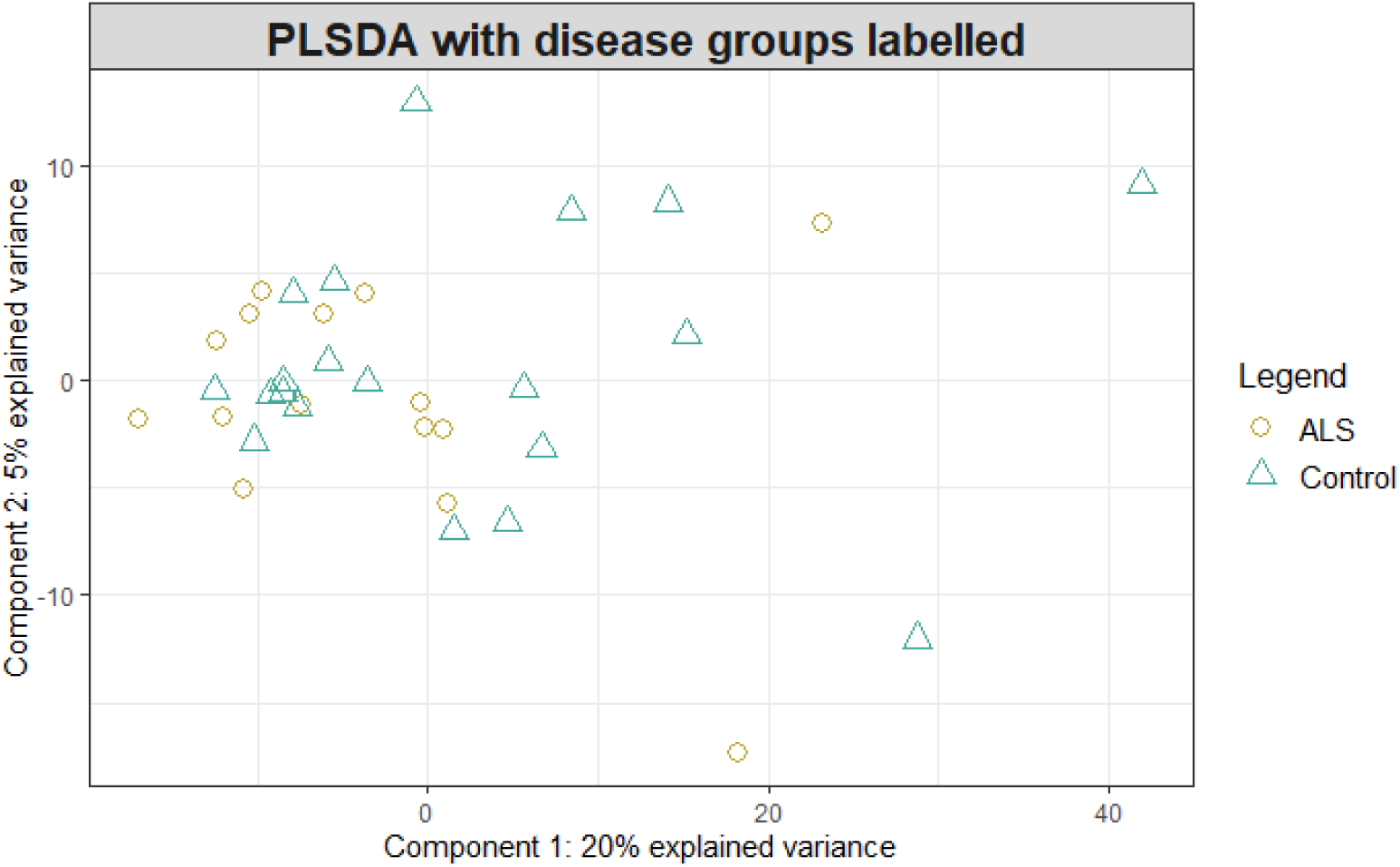
Three component PLSDA model separating TDP-43 and ferritin pathology groups with points labelled by disease status rather than pathology group. The identified altered metabolites explain variance in data dependent on pathology presence but lack explanation of separation between disease vs control.

**Supplementary figure 3.**
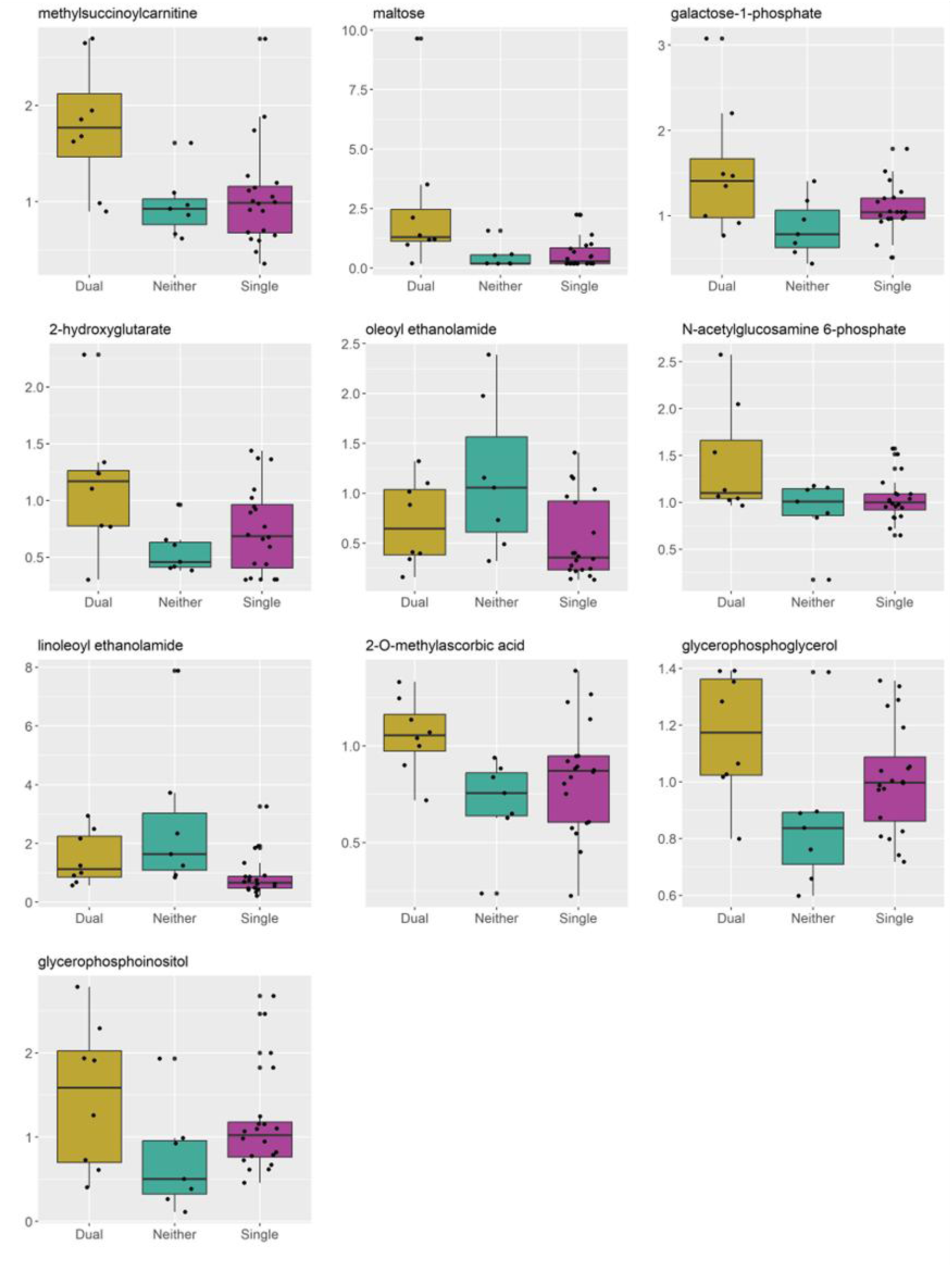
Boxplots to show relationships between metabolites with top 1O VIP scores and pathology status.

**Supplementary figure 4.**
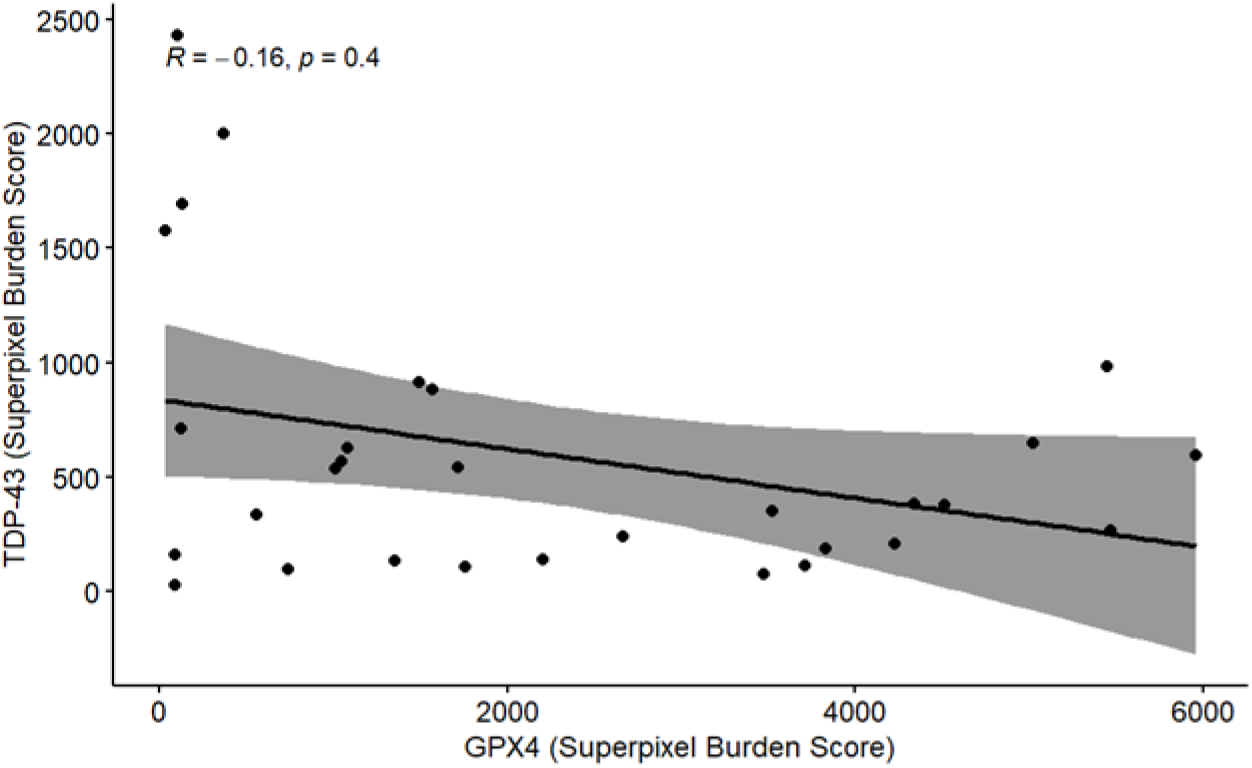
There is no significant correlation between TDP-43 and GPX4 when assessing superpixel burden score (Spearman; R=-0.16; p=0.4).

## Notes

### Competing Interest Statement

The authors have declared no competing interest.

